# Quantifying Platelet Margination in Diabetic Blood Flow

**DOI:** 10.1101/344655

**Authors:** H.-Y. Chang, A. Yazdani, X.J. Li, K. A. A. Douglas, C. S. Mantzoros, G. E. Karniadakis

## Abstract

Patients with type 2 diabetes mellitus (T2DM) develop thrombotic abnormalities strongly associated with cardiovascular diseases. In addition to the changes of numerous coagulation factors such as elevated levels of thrombin and fibrinogen, the abnormal rheological effects of red blood cells (RBCs) and platelets flowing in blood are crucial in platelet adhesion and thrombus formation in T2DM. An important process contributing to the latter is the platelet margination. We employ the dissipative particle dynamics method to seamlessly model cells, plasma, and vessel walls. We perform a systematic study on the RBC and platelet transport in cylindrical vessels by considering different cell shapes, sizes and RBC deformabilities in healthy and T2DM blood, as well as variable flowrates and hematocrit. In particular, we use cellular-level RBC and platelet models with parameters derived from patient-specific data and present a sensitivity study. We find T2DM RBCs, which are less deformable compared to normal RBCs, lower the transport of platelets toward the vessel walls whereas platelets with higher mean volume (often observed in T2DM) lead to enhanced margination. Furthermore, increasing the flowrate or hematocrit enhances platelet margination. We also investigated the effect of platelet shape and observed a non-monotonic variation with the highest near-wall concentration corresponding to platelets with moderate aspect ratio of 0.38. We examine the role of white blood cells (WBCs), whose count is increased notably in T2DM patients. We find that WBC rolling or WBC adhesion tend to decrease platelet margination due to hydrodynamic effects. To the best of our knowledge, such simulations of blood including all blood cells have not been performed before, and our quantitative findings can help separate the effects of hydrodynamic interactions from adhesive interactions, and potentially shed light on the associated pathological processes in T2DM such as increased inflammatory response, platelet activation and adhesion, and ultimately thrombus formation.

## INTRODUCTION

Patients with type 2 diabetes mellitus (T2DM) are at high risk of developing cardiovascular diseases, since T2DM is clearly linked with the onset of a thrombotic event (1, 2). Several factors contribute to the prothrombotic status in T2DM, such as elevated coagulation, impaired fibrinolysis, endothelial dysfunction and platelet hyperreactivity (3–5). Among them, the role of platelet in the formation and development of thrombi is of particular importance. Platelets in T2DM are characterized by dysregulation of several signaling pathways, and they can become hyperreactive and more prone to undesirable activation, adhesion and aggregation (6–9). Other abnormalities such as raised platelet count and high mean platelet volume (MPV) have also been observed in T2DM (10, 11). Shilpi *et al.* (12) showed that MPV (11.3 ± 1.0 fL) in diabetics is significantly higher than the MPV (9.0 ± 0.6 fL) in non-diabetics. For diabetic patients with micro- or macrovascular complications, MPV can be even higher (13, 14). Papanas *et al.* (13) found that T2DM patients without retinopathy have MPV = 10.9 ± 1.1 fL while those with retinopathy have MPV = 15.8 ± 1.3 fL. It is reported that large platelets are more reactive than smaller platelets (15), produce more prothrombotic factors such as thromboxane A2, and express a greater number of adhesion receptors, such as P-selectin and GpIIb/IIIa (16).

The process of thrombus formation and development at an injured site on a blood vessel wall involves a number of simultaneous processes from flow dynamics, blood coagulation cascade and transport of tissue factors (TF), to platelet adhesion, activation, and aggregation (17–19). Thrombosis can occur in the arterial or the venous circulation, and it is triggered by a disruption in endothelial cells and the exposure of TF and collagen to blood. Accelerated atherosclerosis observed in T2DM would increase the risk of endothelial erosion of plaques, followed by thrombus formation (20). For efficient adhesion to an injured site, platelets must be sufficiently close to the site for platelet–subendothelium or platelet–platelet bonds to form. In a shear flow, red blood cells (RBCs) experience a wall-normal force which arises due to their deformation, and propels them away from the vessel wall, leading to formation of a cell-rich region in the core of the vessel and a cell-free layer (CFL) near the wall (21–23). This phenomenon was first quantified by Fahreus and Lindqvist (24). Deformable RBCs, on the other hand, can expel rigid platelets from the core of the stream towards the CFL region. Such transport and accumulation of platelets near the wall known as platelet margination occurs in the presence of RBCs at sufficient hematocrit (*H_ct_*) and flowrate.

Significant progress has been made in understanding the platelet margination probability and dynamics, which characterize the physical contacts of platelet with the vessel wall and its direct or indirect effects on platelet adhesion. The first *in vivo* experiments of platelet distribution in venules and arterioles of rabbit mesentery were performed by the group of Reneman *et al.* (25–27). They found that the normalized platelet density is concentrated near the wall of arterioles, but the opposite platelet distribution is observed in venules. This discrepancy may be due to the rheological effects of RBCs, different position of a vessel in the vascular network, and also vessel wall properties. Numerous *in vitro* studies on the platelet margination and adhesion have also been reported (28, 29). Aarts *et al.* (30) examined the radial distribution of RBCs and platelets in 3-mm cylindrical vessels at various hematocrit and wall shear rates by the laser-Doppler technique. They found that near-wall concentration of platelets increases with increasing hematocrit and wall shear rates. Similar results of enhanced margination of platelet-sized spherical particles for *H_ct_* = 20% compared to for *H_ct_* = 10% were found by Fitzgibbon *et al.* (31). In addition to the experiments, they conducted boundary integral simulations to confirm their experimental results and to track the trajectories of the particles with higher resolution. Eckstein and coworkers (32, 33) performed several studies on the lateral transport of platelet-sized latex beads in flows of blood suspensions. They found that the near-wall excess in concentrations of platelet-sized beads occurred when *H_ct_* >10% and the wall shear rate > 200 s^−1^, while the largest amount of lateral transport of beads was observed at moderate shear rate ~560 s^−1^.

Along with the aforementioned experimental studies, recent advances in computational modeling and simulations enable us to investigate blood flow dynamics under physiological and pathological conditions (34–37). Crowl and Fogelson (38, 39) have performed blood flow simulations in a two-dimensional channel using a Lattice Boltzmann Method. They found that the lateral diffusivity of a platelet is correlated with its lateral position, and platelet margination mostly occurs after the CFL has fully developed. The numerical studies from the group of Aidun *et al.* (40–42) demonstrated that, in addition to the hematocrit, the viscosity ratio of cytoplasm within the RBC to suspending fluid as well as the platelet shape play an important role in the margination process; less viscous cytoplasm of RBC than plasma or more spherical shape of platelets can enhance the platelet margination rate. Vahidkhah *et al.* (43) have captured the three-dimensional nature of RBC-platelet collisions. For platelets approaching to the wall, anisotropic platelet diffusion within CFL leads to the formation of platelet clusters, which might contribute to clot formation. Soares *et al.* (44) and Yazdani *et al.* (45) have performed simulations of platelet transport in complex geometries resembling arterial stenoses. They employed dissipative particle dynamics (DPD) to investigate the transport dynamics and self-orbiting motions of platelets when they flow through a constriction. Yazdani and Karniadakis (45) also found that higher levels of constriction and wall shear rates enhance platelets margination significantly, which may consequently lead to enhanced post-stenosis platelet deposition onto the thrombus. In addition, the effects of microvessel tortuosity and the pathological alteration in the platelet size and density on the platelet activation and thrombus formation have been systematically investigated by Chesnutt and Han (46). In their study, however, they considered platelets as spherical particles and neglected the effects of RBCs for computational efficiency.

Following our previous work on modeling diabetic RBCs (47), here we extend our simulations to complex blood flow in microcirculation in order to study correlations between T2DM and increased platelet margination. We perform high-fidelity computational studies of blood flow through cylindrical vessels, and focus on the radial distribution of diabetic RBCs and platelets, under conditions resembling blood flow in an arteriole for T2DM subjects. In particular, we incorporate the properties of diabetic RBCs and platelets from available experiments and also a new dataset collected from 64 diabetic patients. Several hemodynamic factors such as blood flowrate, hematocrit, and platelet shape are taken into account. In addition, there is growing evidence showing that white blood cells (WBCs) contribute to the disturbances of microcirculation in diabetes due to the promoted inflammatory cell recruitment (48–50). We further consider the influence of WBC dynamics on platelet transport in blood vessels by respectively incorporating leukocyte circulation, rolling along the vessel wall, and firm adhesion to the wall into our simulations. Such *in silico* hematologic study involving large population of blood cells enables us to gain insights in the circulation and distribution of RBCs, platelets, and leukocytes in diabetic conditions, which in turn will help us understand the connection between enhanced thrombosis and inflammation in T2DM (51).

## METHODS

We employ dissipative particle dynamics (DPD) to model whole blood flow, *i.e.* plasma, RBCs, platelets and WBCs, in 40-micron diameter vessels. DPD is a coarse-grained analog of molecular dynamics, where each particle represents a lump of molecules that interacts with other particles through soft pairwise forces. In addition to blood plasma modeled by collections of free DPD particles, the membrane of suspending cells including RBCs, platelets, and WBCs is constructed by a 2D triangulated network with *N_v_* vertices (DPD particles). These vertices are connected by *N*_s_ elastic bonds to impose proper membrane mechanics. More details of hydrodynamic interactions between DPD particles, and models for blood cells are presented in Appendix A. We also validated systematically the DPD model by simulating the margination of spherical particles of different sizes in blood flow and compared against recent experimental measurements (52). The validation results are given in Appendix B.

### Parameter estimation

#### RBC models

Under physiological conditions, a healthy RBC has a large area-to-volume (S/V) ratio with biconcave shape and a remarkable deformability and adaptability in mechanical behavior. In this study, we model a RBC at normal state (NRBC) with *N_v_* = 500, shear modulus *µ*_0_ = 4.73 *µ*N/m and bending rigidity *k*_0_ = 2.4×10^−19^ J. A NRBC is set to have the cell surface area 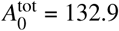, and cell volume 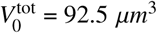, which give S/V = 1.44. All parameters used in our NRBC model are extracted from available measured data and the representative RBC model has been extensively used from single cell mechanics to blood flow dynamics (22, 23, 53, 54). Different from NRBCs, diabetic RBCs possess decreased cell deformability and are known to exhibit shape change from a normal biconcave shape to a near-oblate shape with a smaller S/V ratio (55–57). In addition, it was observed that the increase in mean corpuscular volume (MCV) is related to high blood glucose level (58, 59). Diabetic RBCs with high intracellular glucose will induce osmotic imbalance between RBCs and their surrounding plasma. Morse *et al.* have shown an increase in MCV from 93 to 114 *µm*^3^ by *in vitro* incubation of normal RBCs with glucose concentration of 20 mg/ml for an hour. A further increase in MCV (> 120 *µm*^3^) can be achieved with increasing incubation temperature (59). This MCV change is reversible once the glucose concentration is diluted. Agrawal *et al.* also claimed that the swollen RBCs observed from the sample blood of T2DM patients can be associated to the possible metabolic disturbance (60).

In this work, we have performed an evaluation of several clinical characteristics in blood samples from 136 human subjects, see Table S4. The number of patients diagnosed of T2DM is 64 with a mean age 61 ± 13.7 years compared to 72 patients with a mean age 52 ± 17.8 years in controls. The blood glucose parameters, HbA1c and fasting blood glucose (FBG), are significantly higher in diabetics (HbA1c = 7.66 ± 1.41% and FBG = 144.5 ± 48.9 mg/dl) compared to the controls (HbA1c = 5.25 ± 0.29% and FBG = 99.0 ± 15.4 mg/dl). Details of the blood sample preparation and analysis procedures are included in Appendix C. As shown in Fig. S3(a) and (b), diabetic RBCs exhibit no statistically different MCV compared to that of the normal RBCs. A recent study performed by Lee *et al.* (61) shows a reduction in the membrane fluctuation of diabetic red cells, but there was no statistical difference in volume and surface area distributions of non-diabetic and diabetic RBCs. Although the alteration in the size and shape of diabetic RBCs is still debatable and probably dependent on glycemic control of the study subjects, we considered extreme values for S/V that reported by Jin *et al.* (56). We have recently proposed a T2DM RBC model (47), which has been validated with available experimental data. Following our previous study, here, we extend our T2DM RBC model (DRBC) to a suspension with cell properties similar to those of a NRBC except for a shear modulus, *µ*_s_ = 2*µ*_0_ (*µ*_0_ is the shear modulus of healthy subjects), and a reduced S/V ratio, S/V = 1.04.

#### Platelet models

An inactivated platelet is an oblate spheroid that can be characterized by its aspect ratio (AR) defined as the ratio of the minor axis to major axis. The typical AR range for a normal platelet is 0.25 – 0.5 and the mean platelet volume (MPV) is 6 – 10 fL (*µm*^3^) (62, 63). However, previous studies have revealed a higher MPV in diabetic patients than in non-diabetic controls, which is considered as a risk factor for vascular diseases (12). As shown in our statistical study in Figs. S3(c) and (d), the scattered data points and the number distribution of MPV are significantly higher in the diabetic group than in the control group. The average MPV in diabetic subjects is 10.0 ± 1.26 fL compared to 9.2 ± 1.70 fL in non-diabetics with a *p*-value of 0.003. Such enhancement in the MPV of diabetic platelets may be due to osmotic swelling as a result of hyperglycemia. Patients with less well controlled diabetes are the ones developing micro- and macrovascular complications such as retinopathy. In a previous study by Papanas *et al.* (13), diabetic patients with retinopathy had MPV = 15.8 ± 1.3 fL which corresponds to a ~5 fL difference between the MPV (10.9 ± 1.1 fL) of diabetics without retinopathy. Levels of glycemic control may explain the difference in measured MPV between our and the other studies (11–14). Indeed, this could have significant implications since the effect observed here could be much more pronounced in diabetics with poor glycemic control who by definition are at much higher risk for developing cardiovascular disease (CVD) (1). Large studies with a wide range of HbA1c levels are needed to fully clarify this issue. Another factor not studied here is the potential modifying effect of smoking which has vasoconstrictive effects and is known to have multiplicative effects in terms of increasing CVD risk among diabetics (64).

Considering these facts, here, we started our efforts in this field by using the platelet model with parameters driven from patient-specific data, and further conduct a sensitivity study. A normal platelet (PLT) has cell volume 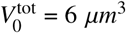 while a diabetic platelet (PLT^*^) has 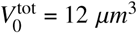. The aspect ratio AR = 0.38, number of vertices in membrane network *N_v_* = 48, and elastic properties of platelets *µ_s_* = 100*µ*_0_ and *k_c_* = 10*k*_0_, are given to all platelet models. Note that platelets are non-deformable in their resting state and thus, we use high values of shear modulus and bending rigidity to impose the proper cell stiffness.

#### WBC model

A WBC is represented as a triangulated sphere with *N_v_* = 2498 and diameter of 10 *µm*, which is similar to the previous leukocyte model used by Lei and Karniadakis (65). Compared with healthy RBCs, a WBC is less deformable with the estimated membrane stiffness 0.3 – 1.2 dyn/cm (66). It is also reported that impaired deformability has been observed in children with diabetes (67), but the changes in WBCs shape are unclear. In this study, we model a WBC with membrane elasticity of *µ_s_* = 22*µ*_0_ and *k_c_* = *k*_0_.

#### Fluid-cell, cell-cell, and fluid (cell)-wall interactions

Blood plasma is modeled by DPD fluid with viscosity *η* = 1.2×10^−3^ Pa·s. The fluid–cell interactions are accounted for through viscous friction using the dissipative and random DPD forces. The repulsive-force coefficient for the coupling interactions is set to zero. To prevent cell-cell overlap, we typically use relatively strong conservative repulsive forces between DPD particles on the membranes of blood cells. Additionally, we employ the Morse potential to ensure strong repulsive forces among cells and prevent their overlap. We refer to our previous work (45) for the details of the DPD model. The parameters used for the fluid–cell and cell–cell interactions are given in Tables S1–S3 in the Appendix A. In this study, we apply the same interaction parameters for both platelets and WBCs. The wall is modeled by frozen DPD particles with the same radial distribution function as the fluid particles in combination with a specular reflection of the plasma and cell particles at the fluid-wall interface (22, 23). The scaling of model and physical units are also identical to those presented in (45). In all simulations, the length scale *r^M^*= 1.0×10^−6^ m and the time scale *τ*= 1.8×10^−4^ s are adopted.

### Simulation setup

The blood flow domain is a cylindrical vessel with diameter D*_t_*= 40 *µm* and periodic in the flow direction as shown in Fig. 1(a). NRBCs in their resting form are biconcave in shape while DRBCs are near-oblate in shape. PLTs and PLT*s are both in oblate spheroid shape with AR = 0.38, but a PLT* has cell size twice than the size of PLT. In this study, the volume fraction of RBCs, *i.e.* average hematocrit (*H*_*ct*_), is varied while the number of platelets in the blood is fixed at 437,000 platelets/mm^3^ (128,000–462,000 platelets/mm^3^ for normal blood (68)). All fluid particles and blood cells are randomly placed within the vessel for the initial condition. To drive the flow (simple fluid or blood) in the vessel, we apply a constant body force to each DPD particle in the flow direction. The simulations are performed for sufficient time to achieve steady state, which is characterized by a time-independent fully developed velocity profile, *i.e.* a plug-like profile for blood. In this study, the time to reach steady state is equal to approximately a half second in physical units. Fig. 1(b) shows the time-averaged axial velocity distribution for the healthy blood at *H_ct_* = 15%. The data is fitted with a parabolic curve (red line) that follows a plug-like characteristic. This plug-like velocity profile is due to the non-uniform distribution of RBCs in the vessel. RBCs in Poiseuille flow migrate to the vessel center forming a depleted region, the cell-free layer (CFL), at the periphery of a vessel. A sample snapshot of NRBCs concentrated in the core region and PLTs marginated toward the wall is shown in Fig. 1(c). To vary the flowrate, we define a characteristic shear rate 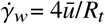, where *ū* is the mean velocity and *R_t_* is the vessel radius. 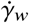 corresponds to the wall shear rate of a parabolic Poiseuille flow with the same *ū*, and is varied in our simulations. In addition, the cell Reynolds number is defined based on 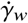, RBC diameter (*D_RBC_*), and the kinematic viscosity of plasma (*ν*), 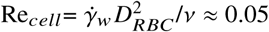, which renders the inertia effect negligible.

**Figure 1:**
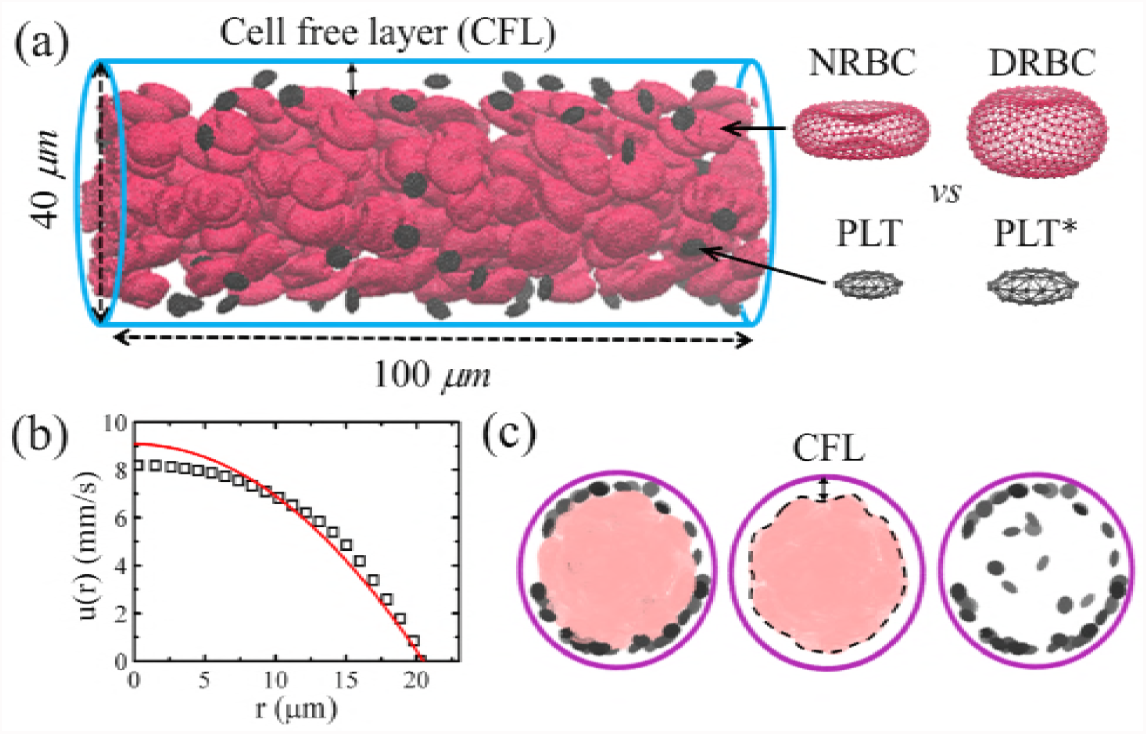
A typical simulation setup. (a) Side view of the healthy blood flow in a vessel with diameter D*_t_*= 40 µ*m* and hematocrit = 15%. Cell free layer (CFL) is evident at fully developed blood flow, red cells are deformable NRBCs with biconcave shape in their resting form and small black cells are PLTs. Diabetic blood in a vessel is composed of DRBCs with near-oblate shape in their resting form and larger in size PLT*. (b) A plug-like velocity profile is achieved and a parabolic curve (red line) is superimposed for reference. (c) Front views of the vessel. Snapshots from left to right show the distributions of red cells and platelets at steady state.

Our clinical data as well as previous studies have shown increased WBC recruitment and count in blood circulation in T2DM (48–50). In diabetics, P-selectin and intercellular adhesion molecule-1 (ICAM-1), which play important role in the adhesion of leukocytes to endothelium, are significantly elevated in human retina and choroid (69). In addition, the rise in WBC count has been observed in our blood samples from diabetic patients (8.04 ± 2.16 *k*/*µL*) compared to control subjects (6.85 ± 1.96 *k*/*µL*), see Table S4. We use our computational framework to investigate three different states of WBC dynamics in blood vessels: circulating in the blood stream, rolling on the wall, and adhered to the wall. For the last two states, we assume that WBCs and endothelial cells are activated and there are adhesive interactions between the receptors of WBCs and the associated ligands expressed on the endothelial surface. Depending on the type and strength of adhesive interactions, WBCs could roll or adhere on the endothelial surface. At physiological shear stress, rolling on selection molecules (*e.g.* P- and E-selectins) are a prerequisite for chemoattractant-stimulated interaction of WBC integrins with endothelium ICAM-1 that arrests rolling WBCs and strengthens adhesion (69–71). To simulate WBC rolling behavior, we initially place WBCs close to the wall and apply a weak conservative repulsive force between DPD particles of WBCs and the wall. To set up blood flow with adherent WBCs, we first run a short period of simulation with rolling WBCs, then freeze those DPD particles located at the interface between WBCs and the wall.

## RESULTS AND DISCUSSION

Having validated our cell transport model with *in vitro* experiments (52) in Appendix B, here we first present the results of healthy blood flow in a vessel and make comparisons with other numerical and experimental studies for further validation. We then study blood flow in diabetic conditions using a suspension of T2DM RBCs (DRBCs) and T2DM platelets with a higher MPV value (PLT*s). The time-averaged concentration profiles of modeled RBCs and platelets flowing through vessels are plotted for different flowrate and hematocrit values. In addition, we investigate the shape effects of platelets on their transport. Subsequently, we take into account the WBCs by including their different dynamics in our flow simulations and analyze their contributions to platelet dynamics and transport.

### Flow of healthy and diabetic blood

Fig. 2 shows the local concentration profiles of platelets and RBCs in the vessel (D*_t_*= 40 *µm*). Here, blood is in two states, i.e., healthy or diabetic, while the wall shear rate and average hematocrit are kept fixed at 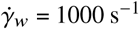 and *H_ct_* = 20%, respectively. All simulations are performed for at least 3000 DPD time units (≈ 0.54 sec) to achieve steady flow conditions, and the last 500 time units are used for the estimation of the time-averaged concentration of RBCs and platelets. These time-averaged concentration profiles are also spatially averaged along the flow direction and plotted against the radial distance from the wall. A histogram bin in the platelet concentration profile represents the cell-center density of platelets within a cylindrical shell (with volume *V_j_*) having a thickness of 0.5 *µm* (platelets/*µm*^3^). The local RBC concentration profile is the cell-center density distribution normalized with respect to the mean prescribed density in that shell (*ρ_j_*). For example, we have 270 NRBCs and hence *ρ_j_* = 270 x *V_j_* /*V_t_*, where *V_t_* is the total volume of the cylindrical vessel; similarly for DRBCs for which we have 200 cells due to their larger size. Therefore, this normalized RBC distribution is adopted to eliminate the cell number difference between the modeled normal (NRBCs) and diabetic red cells (DRBCs).

**Figure 2:**
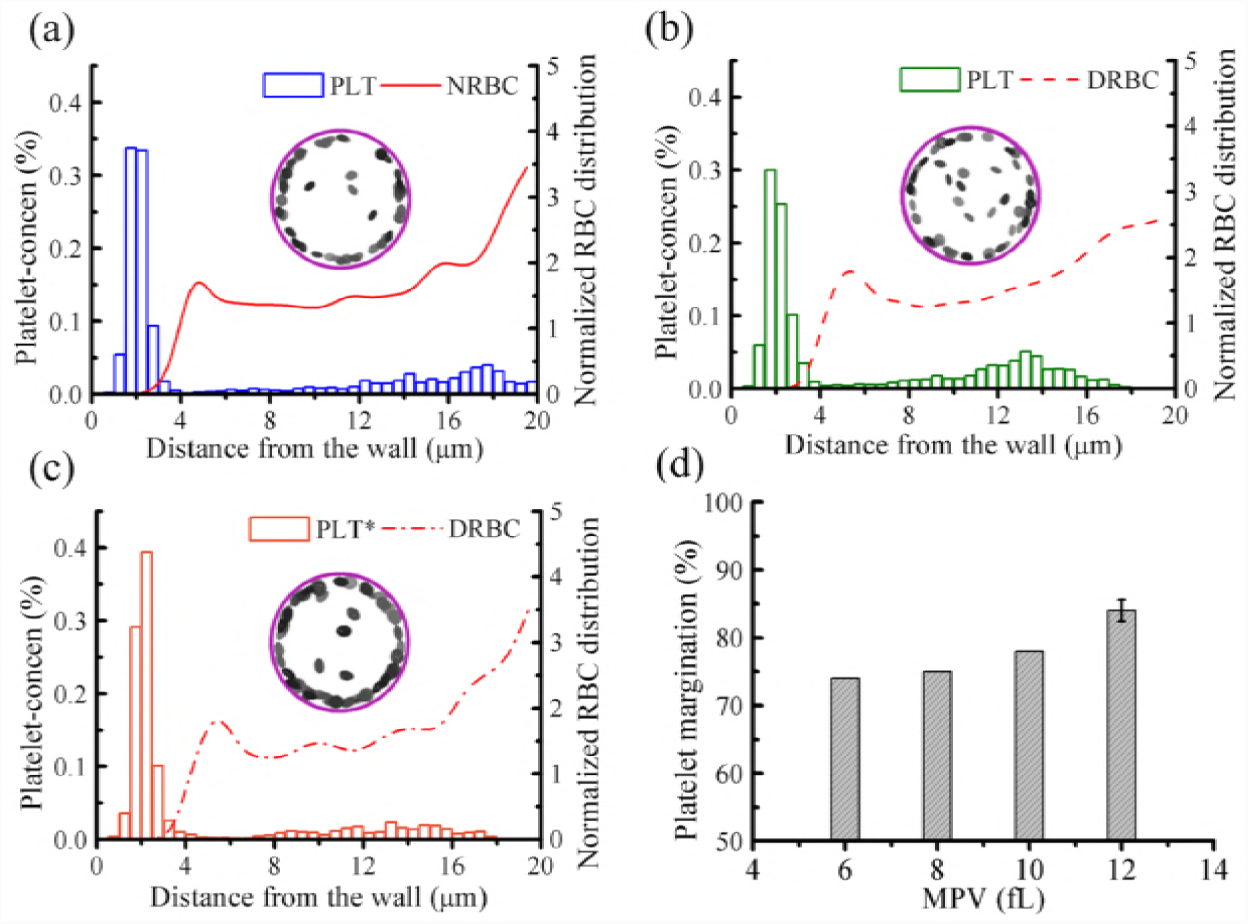
Difference in platelet margination between healthy and diabetic RBC+platelet suspensions. Concentration profiles of platelets and RBCs in the vessel for the cell suspensions of (a) NRBCs+PLTs, (b) DRBCs+PLTs, and (c) DRBCs+PLT*s. Snapshots showing the front views of vessels with platelets only at time= 0.54 sec. (d) Percentage of platelet margination for platelets with different MPV values suspended in diabetic RBCs (DRBCs) and plasma. The vessel diameter D*_t_*= 40 *µm*, average hematocrit *H_ct_* = 20%, and wall shear rate 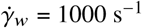 remain the same for all cases.

In healthy blood flow, the distribution of NRBCs is increased around the center of the vessel, exhibiting a primary peak at the centerline and a secondary peak just outside the cell-free region (Fig. 2(a)). Similar RBC distributions were found in previous studies (22, 23). A high cell density of NRBCs in the vessel center is due to the relatively small shear rate in that region, which allows a close packing of NRBCs. Away from the vessel center, the shear rate is increased, leading to the destruction of the close-packed structure of NRBCs. A peak in the distribution occurs next to the CFL, caused by the alignment of NRBCs and their cell-centers that have similar radial positions. On the other hand, non-deformable PLTs are expelled by deformable NRBCs to the wall and dispersed in the CFL region. The formation of the CFL reduces the hydrodynamic resistance and also further promotes platelet margination. The CFL thickness can be determined by *δ* = *R_t_- R_core_*, where *R_core_* is the average radius of the center-occupied RBC core, see also a snapshot of RBC core shown in Fig. 1(c); *δ* ≈ 2.75 *µm* is estimated for the healthy blood with *H_ct_* = 20% flowing through a vessel with diameter= 40*µm*, which is close to that obtained by Mehrabadi *et al.* (41) (*δ* = 3 *µm*) for blood flowing in a channel of height = 40 *µm* and hematocrit = 20%. To quantify platelet margination, we define the percentage of platelet margination (*ϕ_p_*) by the ratio of number of platelets in the cell-free region to the total number of platelets in the simulated blood. A value of *ϕ_p_*= 84% is obtained for the NRBC and PLT suspensions flowing at 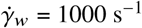.

For diabetic blood flow simulation of DRBC suspensions, a relatively lower platelet concentration in the CFL and *ϕ_p_*= 74% are observed (Fig. 2(b)). The RBC deformability is crucial to the platelet transport, and the impaired deformability of DRBCs compared to NRBCs may cause weaker platelet margination toward the wall. There is *in vitro* evidence showing that the presence of rigid RBCs interferes with the WBC recruitment and platelet margination (72, 73). In addition, Kumar and Graham have presented a detailed analysis of the segregation mechanism incorporating binary mixtures of capsules with different rigidities by performing a computational study with the boundary integral method (74). They found that the degree of segregation increases with increasing rigidity difference between two types of capsules, which is also due to the increase in the heterogenous pair collisions between the stiff and floppy (deformable) capsules. In our study, platelet margination in T2DM RBC suspensions is found to increase when the platelet size is enlarged. As shown in the platelet concentration profiles in Figs. 2(b) and (c), the concentration of PLT* in the CFL region is higher than that of PLT for both cases with identical hematocrit and flowrate. We also conducted a sensitivity study for the platelet transport with different MPV values (MPV= 6 – 12 fL) in the diabetic blood flow, confirming that the percentage of platelet margination increases monotonically with increasing MPV (Fig. 2(d)). The size-dependent margination of particles in blood flow has been found in a number of experimental and computational studies (52, 75–79). For example, it is found that spherical micro-particles (diameter > 0.5 *µ*m) are marginated significantly more than nano-particles (diameter < 0.5 *µ*m), and those small-sized particles are easily trapped in the RBC core. Particle sizes close to the CFL thickness may have the optimized sizes for efficient margination and also for drug delivery systems (80). Comparing the RBC distributions in the vessel, there is no distinct difference between the NRBC and DRBC suspensions as shown in Figs. 2(a)-(c). This result is also in agreement with the studies by Kumar and Graham who showed that an alteration in the membrane rigidity of floppy capsules has minor impact on their distribution (74).

We have also performed simulations of blood flow with a binary mixture of 50% PLTs and 50% PLT*s suspended in DRBCs (Fig. 3). While the normalized cell-center distribution of DRBCs is similar to the previous results, the platelet concentration profile of the mixture (Fig. 3(a)) exhibits mixed features of Figs. 2(b) and (c). To compare the migration velocity of platelets, we plot the trajectories of all modeled PLTs and PLT*s with respect to time, and highlight two representative trajectories. The blue trajectory represents the PLT whereas the red one corresponds to the PLT*. Note that all platelets are initially placed away from the vessel wall with a minimum distance of 5 *µm*. As shown in Fig. 3(b), the larger PLT*s migrate to the wall faster than PLTs, and most marginated platelets show an abrupt lateral displacement once they reach closer to the edge of RBC core. This rapid margination, named “waterfall phenomenon”, was observed in other numerical simulations as well (43, 78). In addition, we estimate the transient percentage of platelet margination of two platelet sizes, see Fig. 3(c). Clearly, there is higher tendency of margination for PLT*s than PLTs at the beginning ~ 0.1 sec, but the difference in the degree of margination becomes smaller at later times.

**Figure 3:**
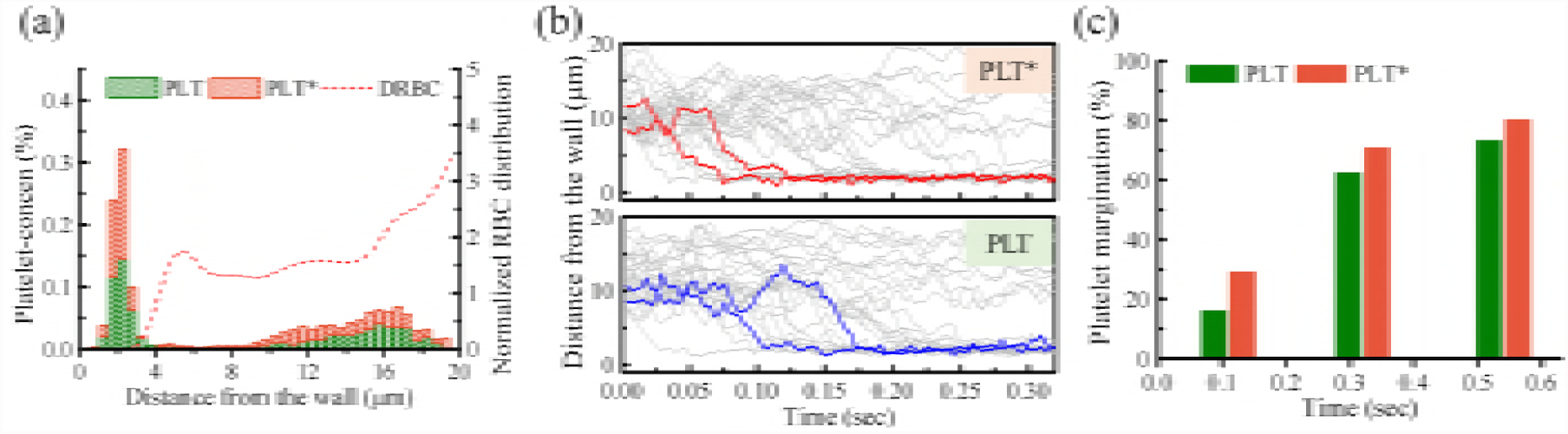
Diabetic red cell suspensions (DRBCs) with a mixture of 50% PLTs and 50% PLT*s flowing in a vessel at 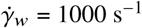 and *H_ct_* = 20%. (a) Concentration profiles of RBCs and platelets, where the histogram plot is adopted for the platelet concentration profile. (b) Trajectories of PLTs and PLT*s with time. Two representative trajectories (the 4^th^ and 10^th^ marginated platelets) are shown in blue lines for PLT and in red lines for PLT*. (c) Temporal evolution of the percentage of PLT and PLT* marginations based on their count.

### Effects of flowrate and blood hematocrit

First, we investigate the effect of flowrate on RBC and platelet distributions in a vessel, where blood hematocrit *H_ct_* = 20% is fixed for three different cell type suspensions: NRBCs+PLTs, DRBCs+PLTs and DRBCs+PLT*s. The RBC distribution and CFL thickness in Figs. 4(a)-(c) seems to be independent of the variation of 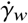 from 1000 s^−1^ (previous case) to 600 s^−1^. This is in agreement with the result of Freund *et al.* (81, 82) that showed the CFL thickness is independent of flowrate when the wall shear rate is high (> 100 s^−1^). Interestingly, although the normalized RBC distributions do not show significant difference at 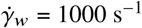 and 600 s^−1^, a decrease in the platelet concentration within the CFL region has been found in all blood cell suspensions. For abetter comparison of the degree of platelet margination in all flow conditions, we estimate *ϕ_p_* (see the values shown in Fig. 4(d)). There is about 6–10% decrease in *ϕ_p_* when wall shear rate is reduced from 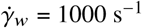 to 600 s^−1^. The decrease in platelet margination with decreasing wall shear rate (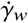) is thought to be the result of reduction in the collision frequencies between RBCs and platelets. In addition, for the case of normal blood flow, NRBCs+PLTs, *ϕ_p_* is close to the diabetic blood flow, DRBCs+PLT*s, and both are higher than the *ϕ_p_* value for cell suspensions containing DRBCs and PLTs. In comparison with the normal platelets, our results indicate that the larger platelets (commonly observed in T2DM) have stronger tendency to migrate toward the vessel wall. This observation together with the platelets hyperreactivity in T2DM could have significant contributions to increased thrombosis events in diabetic subjects.

**Figure 4:**
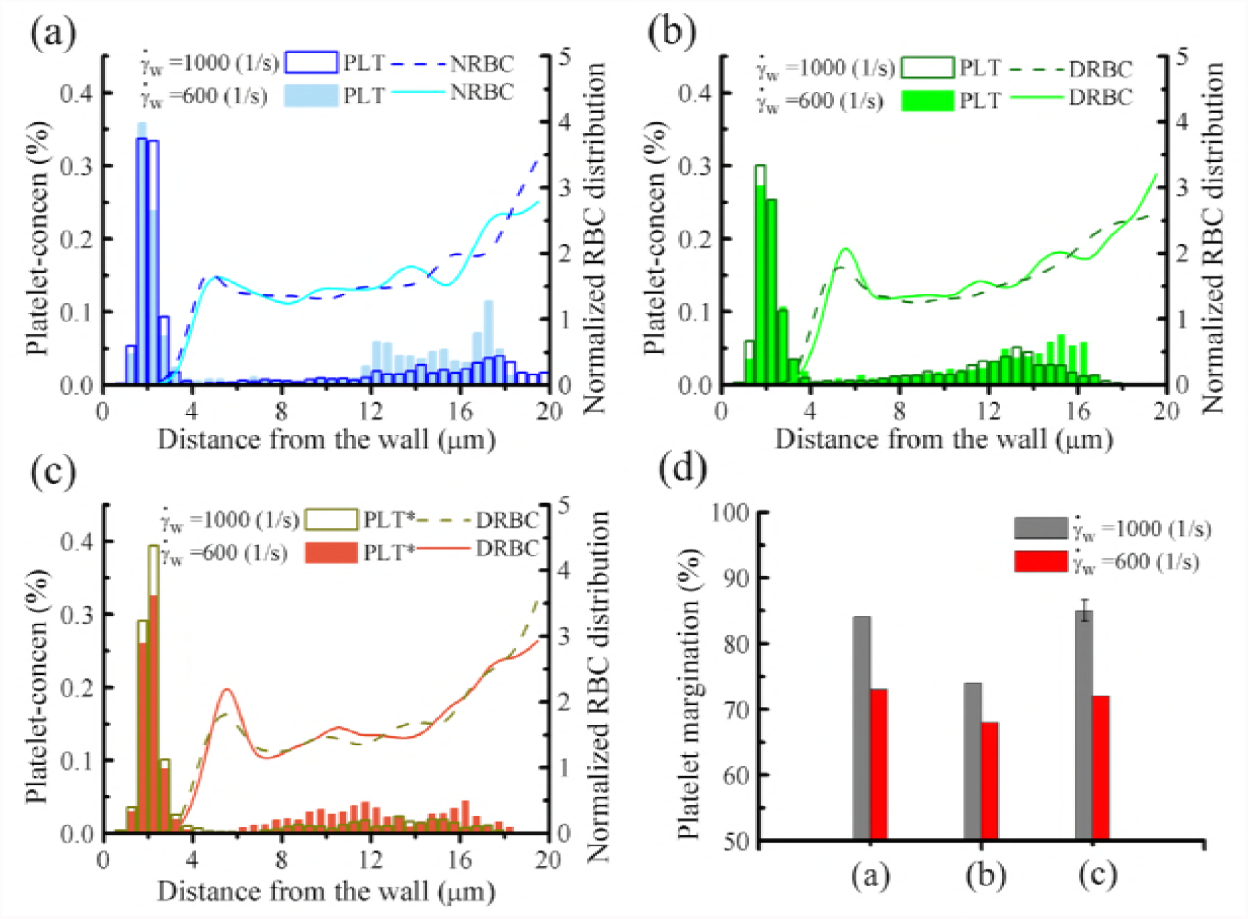
Effect of flowrate. Concentration profiles of platelets and RBCs in the vessel (D*_t_*= 40 *µm*), where cell suspensions of (a) NRBCs+PLTs, (b) DRBCs+PLTs, and (c) DRBCs+PLT*s with identical *H_ct_* = 20% but at different wall shear rates 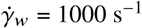 and 1000 s^−1^ are considered. (d) Percentage of marginated platelet *ϕ_p_* for all three cases.

Fig. 5 shows RBC and platelet concentration profiles of normal and diabetic blood flow at different mean blood hematocrit (*H_ct_*), where 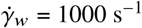. When there is no presence of RBCs (*H_ct_* = 0%), rigid platelets (either PLTs or PLT*s) are concentrated around the center of the vessel (see gray histograms in Figs. 5(a) and (b)). This effect has also been observed in the experiments of particle transport in microchannels without RBCs (52, 83). When RBCs are introduced, the CFL is formed due to the wall-induced migration driving soft RBCs away from the wall, whereas rigid platelets drift toward the wall due to the shear-induced diffusion caused by their heterogeneous collisions with the RBCs. The NRBC and DRBC concentration profiles in Figs. 5(a) and (b) indicate that the RBC core is expanded and CFL thickness is decreased with increasing blood hematocrit *H_ct_*. In addition, as *H_ct_* increases, the near-wall peaks of platelet concentration shift toward the wall and the accumulation of platelets in CFL region is enhanced (see the platelet concentration profiles and *ϕ_p_* estimated for different *H_ct_* in Figs. 5(a)–(c)). We also present the snapshots of diabetic blood flow with mean hematocrit from *H_ct_*= 0% to 25% for a clear visualization of DRBCS and PLT*s distributions in the vessel at time= 0.54 sec (Fig. 5(d)).

**Figure 5:**
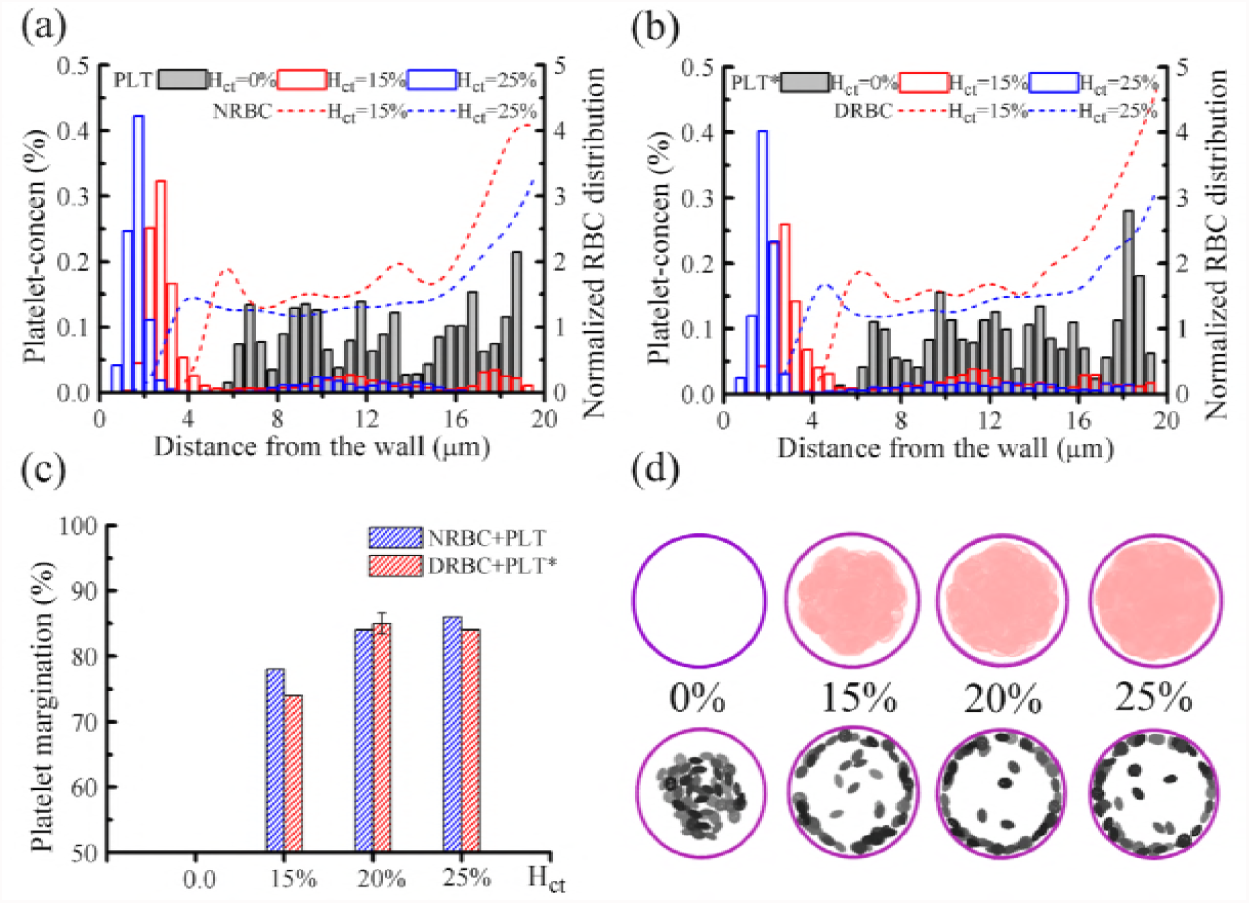
Effect of blood hematocrit at 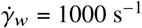. Concentration profiles of platelets and RBCs in the vessel (D*_t_*= 40 *µm*) for (a) *H_ct_*= 0% and NRBC+PLT for *H_ct_*= 15% and 25%; (b) *H_ct_*= 0% and DRBC+PLT* for *H_ct_*= 15% and 25%. (c) Percentage of marginated platelets *ϕ_p_* for normal and diabetic cell suspensions at different *H_ct_*. (d) Front views of the vessel to show the distributions of DRBCs (*upper row*) and PLT*s (*lower row*) with *H_ct_*= 0%, 15%, 20%, and 25% at time= 0.54 sec.

### Effects of platelet shape and WBC count

In general, an inactivated (resting) platelet has an oblate spheroid shape of aspect ratio AR= 0.25 – 0.5 (63). We further perform simulations to study the effects of platelet shape on its margination by varying the aspect ratio AR= 0.28, 0.38 and 0.5. Fig. 6 presents platelet concentration profiles and the percentage of platelet margination of PLT* (MPV= 12 fL) for different AR values suspended in the diabetic red cells (DRBCs). It is clear that the highest near-wall concentration for PLT* is achieved for AR = 0.38. The significantly flatter oblate shapes with AR = 0.28 and the tumid oblate shapes with AR = 0.5 for platelets show lower near-wall accumulations. This result has also been reported recently by Vahidkhah and Bagchi (84) for healthy blood where a higher margination rate was observed for moderately aspherical particles, but not for the highly aspherical ones. In addition, we observe that platelets with AR = 0.28 have less margination propensity with decreasing blood hematocrit (see Fig. 6(b)). Different from the hematocrit effect analysis on the margination of moderate-oblate-shaped PLT*s (see above), flat-oblate-shaped PLT*s are less sensitive to the change of blood hematocrit. Nevertheless, many studies have found higher margination and binding probabilities for particles with aspherical shape than those with spherical shape (85–87). Platelets with their naturally oblate spheroid shape are endowed with larger contact area compared to the spherical particles of the same volume.

**Figure 6:**
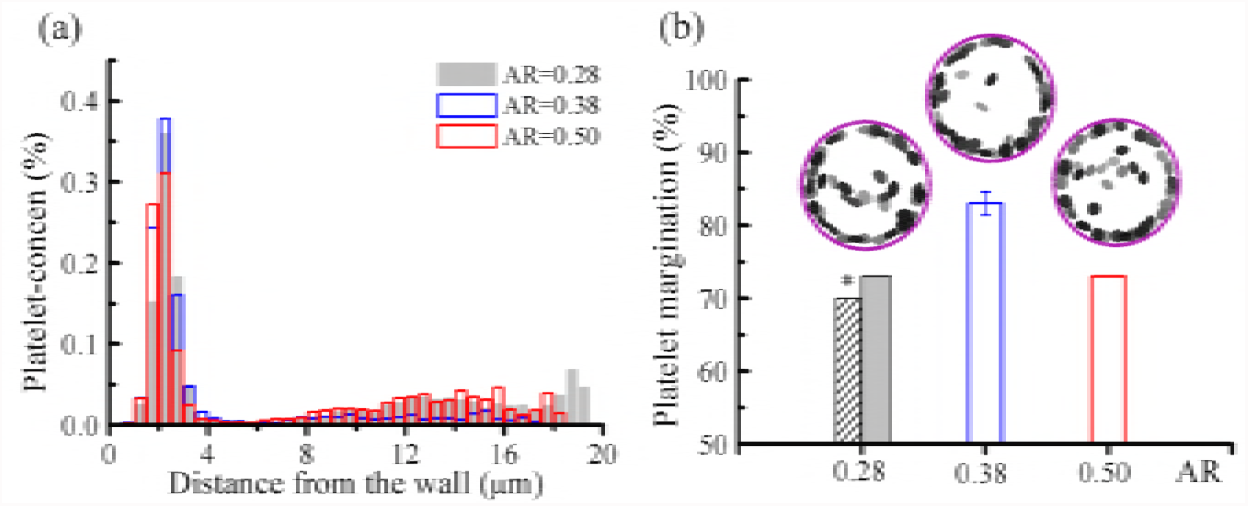
Effect of platelet shape with 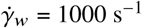 and *H_ct_*= 20%. (a) Concentration profiles of platelets in a vessel with diabetic red cells (DRBCs) and PLT* at different AR values. (b) Percentage of marginated platelets *ϕ_p_* for each AR value. The histogram with * is the result for platelets with AR= 0.28 and blood hematocrit *H_ct_*= 15%. Snapshots show the front views of vessel with different PLT*s suspended in DRBCs, where *H_ct_*= 20%.

White blood cells (WBCs), as an important part of immune system, are subject to a recruitment process from the blood stream to a site of inflammation, followed by a number of interactions with endothelium via receptor-ligand bindings (69–71). For weak or few interactions, WBCs roll along the endothelium, while for strong interactions such as the integrin mediated bonds, WBCs can be firmly attached on the endothelial surface (88). Similar to platelets, WBCs are substantially stiffer than RBCs and their margination occur due to the heterogeneous collisions with RBCs. WBCs margination is prominent in post-capillary venules with a characteristic of low blood flowrate in contrast to the massive number of marginated platelets extensively observed in arterioles (28). However, induced leukocyte-arterial endothelium interactions and the WBC recruitments into atherosclerotic lesions in arteries have been revealed by *ex vivo* experiments (89, 90).

Considering that the blood flow disturbance in small vessels may be caused by the presence of WBCs, we therefore include them into the simulations having three dynamic states often observed in blood vessels: circulating freely in the blood stream, rolling on the wall, and firm adhesion to the wall. The platelet concentration profiles and the percentage of platelet margination in Figs.7(a) and (b) show no clear difference in the near-wall concentration of PLT*s between the blood flows with freely flowing WBCs and without the presence of WBCs. However, we observe that platelet margination is impaired when WBCs emerge near the vessel wall, where more reduction in platelets margination is observed with more rolling or adherent WBCs close to the wall. The solid bars in Fig. 7(b) correspond to a single WBC, whereas empty bars correspond to three WBCs in the domain. We also present three snapshots of near-wall platelet dynamics when platelets transport through or close to an adherent WBC, see Fig. 7(c) and Movie S1–S3. For the platelets encountering an adherent WBC, they will either slide close to the top or the side of a WBC, after which they undergo a flipping motion (Movie S1 and S2). Platelets near an adherent WBC only exhibit a flipping motion in the direction of flow due to their discoid shape (Movie S3). In a flow system with adherent WBCs, platelet transport toward vessel wall is hindered because of the flow resistance induced by the adherent WBCs. However, *in vivo* the cell-cell contacts between platelets and WBCs might be enhanced due to the shear-induced sliding motion of platelets crossing through an adherent WBC. The sliding motion of platelets can facilitate the adhesive interactions between platelet CD62 and leukocyte P-selectin glycoprotein ligand-1 (PSGL-1), which further leads to platelet-WBC aggregations (91, 92). Here we consider only hydrodynamic cell-cell interactions and we will study the effects of adhesive dynamics in future work.

**Figure 7:**
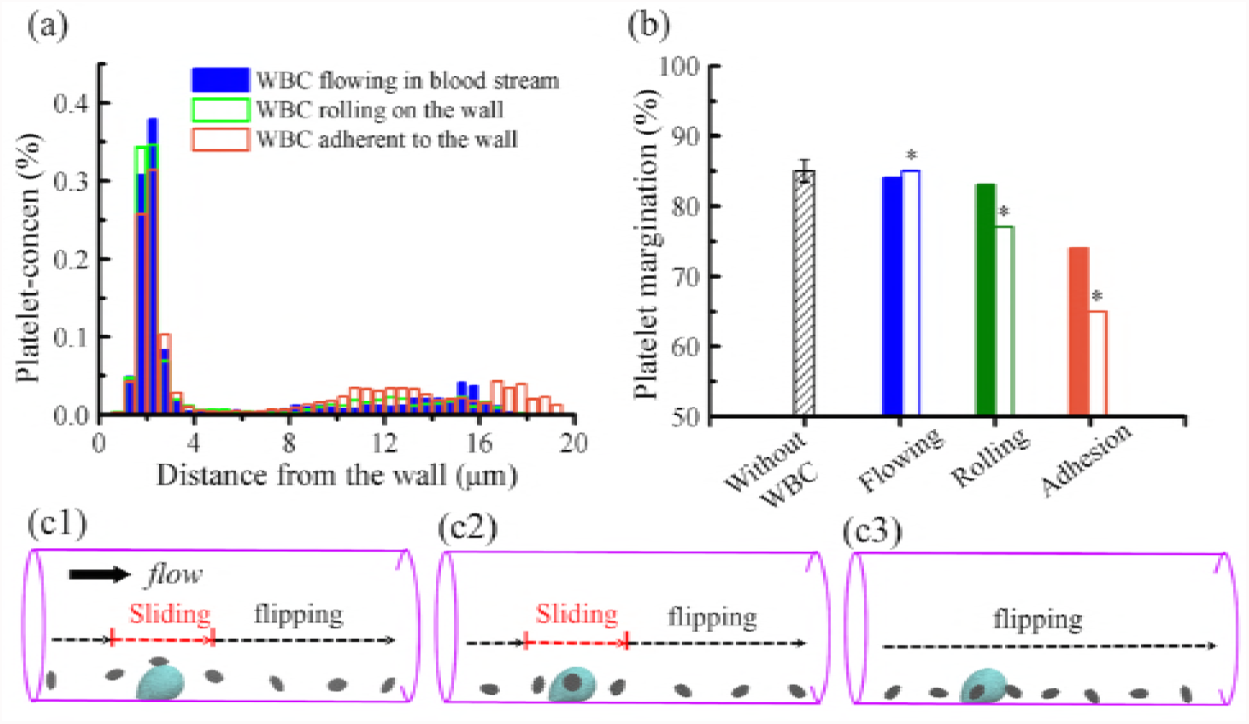
Effect of WBC dynamics at 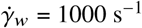 and *H_ct_*= 20%. (a) Concentration profiles of platelets in the vessel with a WBC flowing in the blood stream, rolling on the wall, and adhered to the wall. (b) Percentage of marginated platelets *ϕ_p_* in the blood flow without WBCs and with a WBC under different dynamical states, where * label on the histograms denotes the blood contains three WBCs. (c) Snapshots of near-wall platelet dynamics in a blood flow condition with an adhered WBC showing a representative platelet (c1) sliding from the top of the WBC, (C2) sliding from the side of the WBC, (c3) moving nearby the WBC and flipped by the flow. Diabetic RBCs (DRBCs) and platelets (PLT*) are adopted in these simulations.

## CONCLUSION

Type 2 diabetes mellitus (T2DM) is strongly associated with the accelerated development of atherothrombosis and cardiovascular diseases (1). Thrombus formation is a multi-step process involving platelet transport and the accumulation of tissue factors (TF) at the injured sites. While the elevated levels of TF in T2DM facilitate a cascade of enzymatic reactions, T2DM platelets with enlarged size releasing more vasoactive molecules promote platelet adhesion, activation, and aggregation (12, 13). T2DM RBCs and platelets exhibit abnormal biomechanical properties and biorheology, which can further affect the blood flow dynamics and blood cell transport.

In this study, we perform high-fidelity blood flow simulations to investigate the RBC and platelet transport in the cylindrical vessels resembling small arterioles under healthy and diabetic conditions. In particular, diabetic blood is composed of T2DM RBCs and T2DM platelets. We do not model cell-cell or cell-wall adhesive interactions in order to study systematically the hydrodynamic interactions that induce margination. We employ the dissipative particle dynamics (DPD) method to seamlessly model RBCs, platelets, WBCs, plasma, and the vessel wall. We first validate the DPD model using experimental data for margination of micro-beads (three different sizes) in channel blood flow by Carboni *et al.* (52). We found that, in agreement with the experiments, margination of micro-beads increases with size and shear rate. We have introduced and validated a T2DM RBC model in our previous work (47). Here, we adopted the same red cell model for modeling the diabetic blood, whereas a T2DM platelet model is built based on the parameters informed by the clinical blood analysis of 64 diabetic patients. According to the statistical analysis for the blood characteristics, the average mean platelet volume (MPV) in diabetics (10.0 ± 1.26 fL) is higher than the average MPV (9.2 ± 1.70 fL) in non-diabetics. To distinguish the platelet margination in diabetic blood from that in normal blood, we chose the extreme values of MPV = 6 fL for normal platelets and MPV = 12 fL for T2DM platelets. Similarly for RBCs, we considered their volume (MCV) to have values close to mean value for normal blood and extreme value for diabetic blood. Our blood flow simulations show that platelet margination in whole blood is a very complex and multi-factorial process, affected by the biomechanical properties of cells, their shape and size, as well as flowrate and hematocrit. For diabetic blood flows, we found two competing effects on platelet margination. T2DM RBCs with less deformability compared to normal RBCs contribute to reducing heterogenous collisions, hence impairing platelet margination. On the other hand, T2DM platelets with higher MPV lead to higher margination. We also conducted sensitivity studies for the platelet transport with different MPV values in the diabetic blood flow, and found that the percentage of platelet margination increases monotonically with increasing MPV. Similar to healthy blood flow, platelet margination in diabetic blood flow is increased with increasing flowrate and blood hematocrit. Considering the effect of platelet shape on its margination, we found that the highest near-wall accumulation of T2DM platelets are those with moderate aspect ratio. We further took into account the white blood cell (WBC) dynamics in blood and found that WBC rolling and WBC adhesion tend to decrease platelet margination based purely on hydrodynamic interactions. In future work we will include the effects of cell-cell adhesive interactions as well as cell-wall interactions, which could change some of the quantitative findings of the present work. Elucidating the predictors of platelet margination and of the highest near-wall accumulation of platelets in diabetics will not only allow for the creation of algorithms predicting CVD risk among diabetics but could also point to specific molecular pathways that could lend themselves to predictive or therapeutic interventions in diabetics.

## AUTHOR CONTRIBUTIONS

HYC, AY, XL, CSM, and GEK conceived and designed the research. HYC and AY carried out all simulations. KAAD provided clinical data. HYC, AY, XL, CSM and GEK analyzed the data. HYC, AY, XL, CSM, and GEK wrote the article.

## ACKNOWLEDGMENTS

The work described in this article was supported by National Institutes of Health (NIH) grants U01HL114476 and U01HL116323. CSM also acknowledges support from National Institute of Diabetes and Digestive and Kidney Diseases (NIH/NIDDK) grant K24DK081913. Computations were performed using resources and services at the Center for Computation and Visualization (CCV) at Brown University, the OLCF/ALCF computational resources at ORNL/ANL through an ALCC award, and the NSF-XSEDE resources through award No. TG-DMS140007. HYC would like to thank Dr. Yu-Hang Tang for using an in-house developed code Ermine, and Dr. Zhen Li for the code of cell-generator using in this study.

